# Optimization of mouse embryonic stem cell culture for organoid and chimeric mice production

**DOI:** 10.1101/2020.03.13.990135

**Authors:** Cécilie Martin-Lemaitre, Yara Alcheikh, Ronald Naumann, Alf Honigmann

## Abstract

*In vitro* stem cell culture is demanding in terms of manpower and media supplements. In recent years, new protocols have been developed to expand pluripotent embryonic stem cells in suspension culture, which greatly simplifies cell handling and scalability. However, it is still unclear how suspension culture protocols with different supplements affect pluripotency, cell homogeneity and cell differentiation compared to established adherent culture methods. Here we tested four different culture conditions for mouse embryonic stem cells (mESC) and quantified chimerism and germ line transmission as well as *in vitro* differentiation into three-dimensional neuro-epithelia. We found that suspension culture supplemented with CHIR99021/LIF offers the best compromise between culturing effort, robust pluripotency and cell homogeneity. Our work provides a guideline for simplifying mESC culture and should encourage more cell biology labs to use stem cell-based organoids as model systems.

## Introduction

Maintenance and genetic engineering of pluripotent stem cells in cell culture is a key tool for biomedical and basic research (Evans and Kaufman, 1981; Martin, 1981; Thomson, 1998). For example, production of transgenic mice is efficiently achieved by introducing genetic modifications in ESCs which are modified *in vitro* and are then injected into recipient blastocysts producing chimeric animals (Bradley et al., 1984). In addition, the ability to maintain pluripotent stem cells in culture has fueled the development of differentiation protocols to generate and study various cell types under defined conditions in cell culture (Keller, 1995; Williams et al., 2012). Over the last decade, development of organotypic stem cell differentiation protocols in biomimetic 3D environments has enabled the growth of complex tissues, termed ‘organoids’, which in many aspects can recapitulate the organization and function of organs *in vitro* (Huch and Koo, 2015; Lancaster and Knoblich, 2014; Simian and Bissell, 2017). Organoids provide exciting opportunities for biomedical research, because human diseases can be studied and treatments can be tested *in vitro*, reducing the need for animal models. In addition, organoids will be valuable to study molecular and cellular self-organization rules of mammalian and in particular human-specific morphogenesis.

We recently started to investigate the early morphogenetic events of neural tube formation in synthetic 3D environments. Therefore, we adopted a 3D culture protocol that starts with single mouse embryonic stem cells (mESC) embedded in Matrigel, which self-organize into patterned neural tubes within 6-9 days (Meinhardt et al., 2014; Ranga et al., 2016). This system offers precise control over the initial starting conditions and in principle allows to study the morphogenetic process with high resolution optical microscopy.

To establish the required mESC culture in our lab, we adopted a widely-used culturing protocol which keeps mouse preimplantation embryoblast cells in a self-renewable pluripotent state. The protocol relies on leukemia inhibitory factor (LIF) (Smith et al., 1988; Williams et al., 1988) and cell culture serum to maintain feeder-free adherent mES cells (Nichols et al., 1990; Pease et al., 1990). There are, however, three critical points which make this protocol demanding in terms of time and manpower investment. First, to prevent unwanted differentiation, cells have to be split and transferred into new medium every two days. Second, the passage number has to be kept low requiring frequent reset of the culture. Third, serum is an undefined supplement with varying quality, therefore each new batches of serum have to be rigorously tested. Together, these points make the serum/LIF culture protocol significantly more demanding than standard culture of transformed cell lines, which is part of the notorious reputation that stem cell culture is difficult and expensive.

To simplify mESC maintenance we tested protocols that replace the need for serum by a combination of two kinase inhibitors, specifically MEK inhibitor PD0325901 and GSK3 inhibitor CHIR99021 (2i) (Ying et al., 2008) and/or growth factors (BMP4) (Ying et al., 2003) and combined this with a suspension culture approach (Andang et al., 2008). ESCs cultured in serum-free 2i/LIF medium have homogeneous expression of pluripotency factors and lower lineage-associated gene expression compared to serum/LIF conditions (Marks et al., 2012; Morgani et al., 2017; Wray et al., 2010). Markedly, 2i/LIF cells appear similar to *in vivo* E4.5 epiblast cells on the transcriptome level (Boroviak et al., 2014). However, long-term maintenance of mESC pluripotency by 2i/LIF has been shown to impair the developmental potential of embryonic stem cells via genetic and epigenetic aberrations due to the MEK inhibitor PD0325901 (Choi et al., 2017; Yagi et al., 2017). Suspension culture avoids integrin mediated adhesion of cells to artificial surfaces, which has been shown to prevent unwanted differentiation allowing for longer maintenance of the culture with less splitting effort (Hayashi et al., 2007). In addition, the scalability of suspension culture is higher compared to 2D adherent culture methods (Cormier et al., 2006). While both 2i/LIF and suspension culture have been independently used to simplify mESC culture, the aim of this study was to test if a combination of both protocols could further improve simplicity, robustness and affordability of mESC maintenance.

Here, we tested three suspension protocols using 2i/LIF, 1i/LIF (CHIR99021 inhibitor alone) or BMP4/LIF as self-renewal promoting factors and compared them with the standard adherent serum/LIF protocol. We measured proliferation rates of the culture, germ line transmission in mice, and differentiation into neural tube organoids. We found that 1i/LIF offers the best compromise between reduced culturing effort, robust pluripotency and *in vitro* differentiation.

## Results

We used an R1 mES cell line for all experiments. The cell line was expanded using standard adherent culture supplemented with serum/LIF (Nichols et al., 1990; Pease et al., 1990). To establish suspension cultures we adapted a protocol developed by Andäng et al (Andang et al., 2008). The original protocol uses bFGF/LIF as self-renewal promoting factors. After initial verification of this protocol, we aimed to replace bFGF with 2i/LIF, since 2i has been shown to induce a more homogenous mESC population and to clean-up the culture by eliminating partially-differentiated cells (Marks et al., 2012; Tamm et al., 2013). However, because prolonged MEK inhibition via PD0325901 has recently also been shown to cause significant changes of the DNA methylation state of mESC which compromises developmental potential (Choi et al., 2017; Yagi et al., 2017), we decided to test if mESC can be maintained in suspension without PD0325901 (1i/LIF). In addition, we used BMP4/LIF as a third often used self-renewal supplement. All our suspension culture tests were based on serum-free N2B27 basal medium, similar to 2i/LIF culture protocols (Ying et al., 2008).

### Suspension culture with 1i/LIF promotes self-renewal of mESCs

R1 mESCs were cultured in suspension in N2B27 medium supplemented either with 2i/LIF, 1i/LIF or BMP4/LIF (see *Experimental Procedures*). As a reference we cultured mESCs adherently in serum/LIF. Suspension cells were passaged every 7 days, while adherent cells had to be passaged every 2-3 days. Hence, suspension culture greatly reduced hands-on time for cell maintenance and provided more time to do experiments with cells from the running culture (Figure 1A). As reported before, in suspension, cells formed free-floating spheroids while adherent cells formed 2D colonies (Figure 1B). To quantify proliferation (self-renewal) we determined the cell number after each passage. We found that mESC proliferated under all tested conditions, with 1i/LIF having the highest proliferation rate P = 1.9 million cells/day, while 2i/LIF and BMP4/LIF had proliferation rate P = 1.4-1.5 million cells/day (Figure 1C). Under adherent conditions, proliferation was intermediate with P = 1.6 million cells/day, albeit requiring a higher splitting effort. In addition, adherent culture is typically not kept longer than 4-6 weeks in culture (≈20 passages), because the cells begin to lose pluripotency and the population becomes heterogeneous (Czechanski et al., 2014). This is observed by changes in morphology from dome-shaped to flattened colonies in culture. Furthermore, with higher passage numbers (25 passages, >8 weeks), ES cells have been shown to lose their potential of germline transmission (Longo et al., 1997).

**Figure 1:**
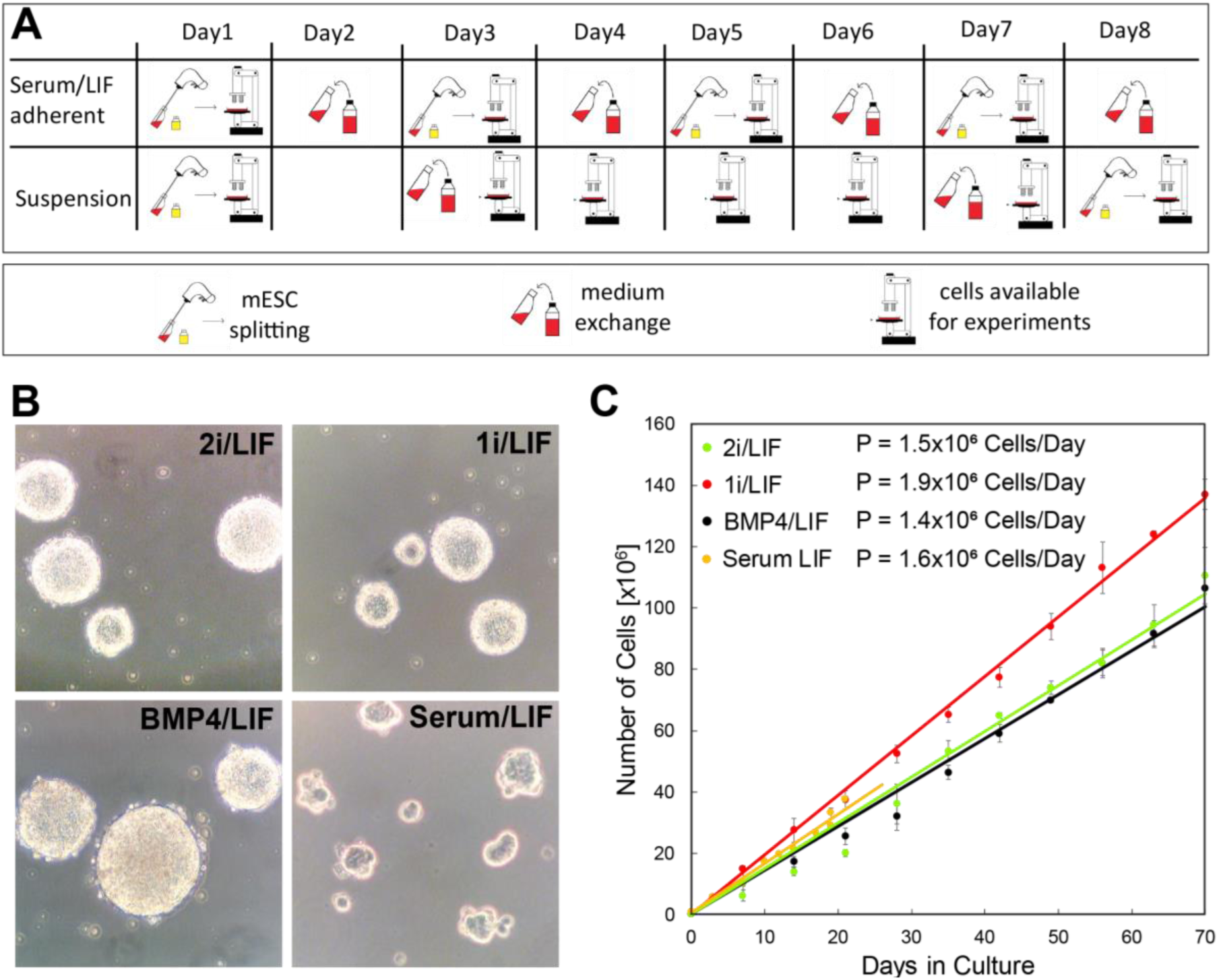
Proliferation of mESC in suspension culture. (A) Scheme for mESC culture handling in suspension and adherent. (B) Bright field images showing morphology of mESC spheroids in suspension (2i, 1i, BMP4) and adherent colonies (Serum). (C) Proliferation rates of the tested conditions. Error is SEM of three independent cultures. Lines show linear fits. The slopes provide the proliferation rates P.

### 1i/LIF suspension mESC can efficiently generate transgenic mice

To test if suspension-cultured mESC were pluripotent, we injected R1 mESC in embryos of Bl6 mice and quantified chimerism and germline transmission. Chimerism was judged by fur color (Figure 2A). Germline transmission was tested by PCR analysis of short tandem repeats (STR) analysis of the sperm of male chimeric animals. In our hands, 1i/LIF suspension cells injected in BL6 mice showed good chimerism rates with high germline transmission efficiency. Even after passages, i.e. 100 days in suspension culture, chimerism and germline transmission were high (≈ 60%) and reached the same efficiency as low passage serum/LIF cells. In contrast, BMP4/LIF suspension cells had a low chimerism rate (< 10%). We also injected 2i/LIF suspension cells at late passage (20 weeks), but were unable to get any chimeric animals.

**Figure 2:**
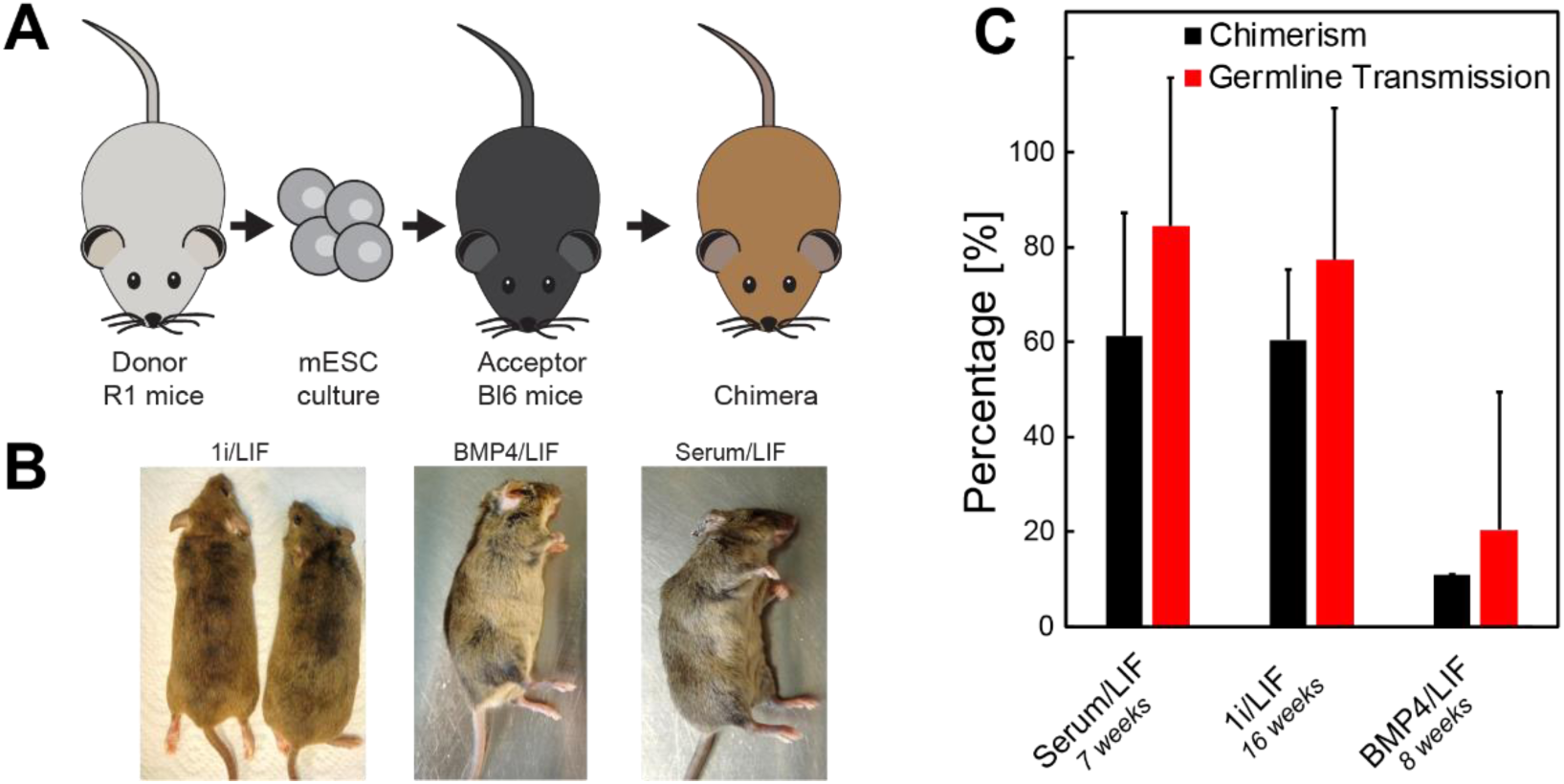
Chimerism and germline transmission of mouse injected mESC. (A) Scheme of mESC injection experiment to test chimerism and germ line transmission (B) Lightening of fur color of R1 injected mESC into BL6 mice indicates chimerism. (B) Quantification of chimerism rate and germline transmission efficiency for mESC cultured in: Serum/LIF (n = 41), 1i/LIF (n = 26), BMP4/LIF (n = 20). Errors are SD.

### 1i/LIF suspension mESC produce neural-epithelial cysts in 3D cell culture

After verifying that mESC efficiently self-renew in suspension culture and can generate transgenic mice with germline transmission, we characterized the potential of these cells to produce neuro-epithelial cyst in 3D cell culture (Meinhardt et al., 2014). We produced single cell suspensions from the four stem cell culture conditions (2i/LIF, 1i/LIF, BMP4/LIF, serum/LIF) and seeded cells in droplets of 70% Matrigel containing N2B27 medium. We then analyzed cell morphology and cell differentiation as a function of time using fluorescence microscopy and flow cytometry (Figure 3).

**Figure 3:**
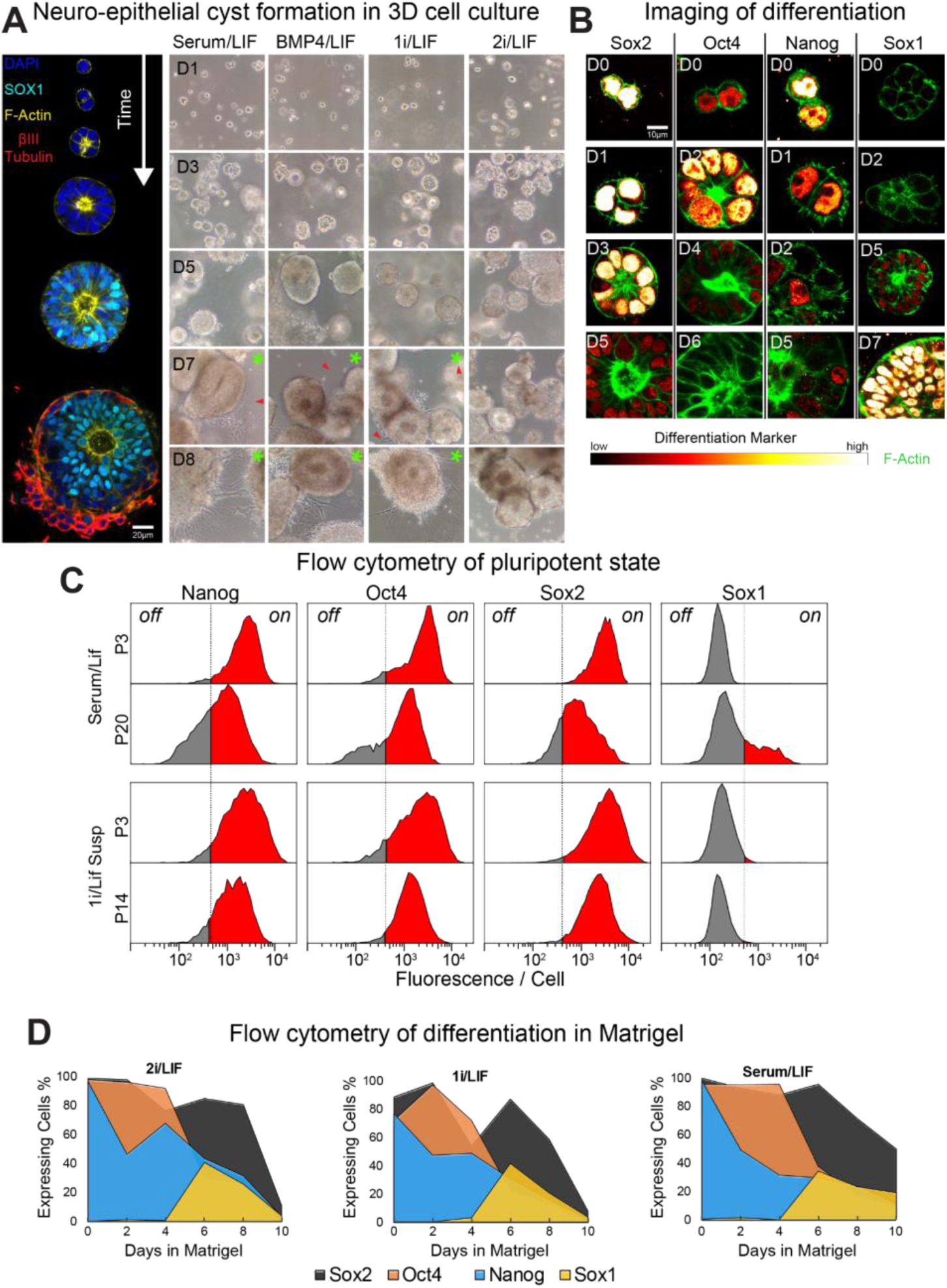
Formation of neuro-epithelial cysts from suspension-cultured mESC in 3D cell culture. (A) 3D Neural-epithelium formation in Matrigel over time. Left, representative mid-sections of confocal images 3D cysts starting from a single cell. After 3-4 days cells form epithelia cysts with a central lumen (actin). Then cells differentiate into neuronal progenitors (Sox1) and finally mature neurons grow at the basal side of cysts (βIII-tubulin). Right, bright field images of the 3D culture started from the different stem-cell culture protocols. Onset of neurogenesis is indicated by green asterisks; red arrows show neuronal protrusion. (B) Fluorescence imaging of differentiation factors after immune-staining of cells from 1i/LIF culture indicates a quick loss of pluripotency (Sox1, Oct4) with the subsequent entry into neuro-ectoderm (Sox1). (C) Flow cytometry analysis of the pluripotent state of mESC cultured in Serum/LIF and 1i/LIF at an early and late passage (D) Quantification of differentiation process in 3D culture by flow cytometry. Day 0 corresponds to the stem cell culture condition. For all time points cells were dissociated fixed and stained with respective marker.

In line with previous reports we found that R1 mESC, which were expanded in serum/LIF, developed into neuro-epithelial cysts within 7 days after seeding in Matrigel with neural induction medium (Meinhardt et al., 2014). During this process cells exist pluripotency and polarize to form an internal apical lumen, then induce neuronal progenitor genes (Sox1) and finally differentiate into neurons at the basal side of the cyst (Figure 3A left). Both lumen opening and formation of neurons could be observed by simple bright field microscopy of 3D cell culture (Figure 3A right panel). We found that cyst shape, lumen formation and onset of neurogenesis of BMP4/LIF and 1i/LIF expanded cells was equal to serum/LIF cultured mESC. Interestingly, mESC that had been expanded in 2i/LIF, showed a two-day delay in neurogenesis.

Next, we aimed to quantify differentiation, i.e. the exit of pluripotency and the entry into neuro-ectoderm, more precisely. We attempted to do this by immune-staining of differentiation markers (Nanog, Oct4, Sox2 and Sox1) at different time-points of the 3D cell culture (Figure 3B). While this strategy in principle worked, i.e. we see the downregulation of pluripotency factor over time, we found that quantification of immuno-stainings over time was very challenging due to technical limitations such as antibody accessibility of the growing tissue. To overcome this limitation, we dissociated cells from the 3D tissue culture at different time points. We then fixed the single cell suspensions, stained for the differentiation markers and performed flow cytometry analysis (FACS). To determine the background signal of cells not expressing the respective marker (*off-cells*), we used control stainings with secondary antibody only (Figure 3C dashed line). We first compared the expression profiles under pluripotent conditions (serum/LIF and 1i/LIF) at early and late passages respectively (Figure 3C). The comparison of the expression histograms clearly shows that, while serum/LIF cells show strong and homogenous expression of pluripotency markers at early passage (P3, 8 days), the population becomes heterogeneous at later passages (P20,50 days) with many cells losing expression of pluripotency factors and inducting neuronal fates (Sox1). In contrast, cells expanded in 1i/LIF suspension show robust expression of pluripotency marker even at later passages (P14, 100 days) with no signs of differentiation into neuronal progenitors (Sox1). Finally, we quantified the timing of exit of pluripotency and entry into neuro-ectoderm by plotting the percentage of *on-cells* for the respective markers over time (Figure 3D). The temporal profiles show that the only obvious difference between the markers tested was prolonged expression of Nanog in 2i/LIF expanded cells compared to 1i/LIF and serum/LIF (Figure 3C). After 4 days in Matrigel culture, 65% of 2i/LIF cells still expressed Nanog, while it was 50% for 1i/LIF and 30% for serum/LIF-expanded cells. This indicates that 2i/LIF-expanded cells take significantly longer to downregulate pluripotency genes, which may explain why these cells show low chimerism rates after mouse injection and delayed neurogenesis in 3D cell culture.

## Discussion

The aim of this work was to optimize and simplify maintenance of pluripotent mES cell culture for the production of transgenic mice and organotypic differentiation in 3D tissue culture. Our results show that suspension culture supplemented with 1i/LIF (CHIR99021) provides a minimal system to expand mES in the pluripotent state over many month, while maintaining the ability to produce chimeric mice and mES cells for *in vitro* differentiation experiments.

By modifying established suspension culture suspension protocols (Andang et al., 2008), we found that CHIR99021/LIF suspension culture condition is sufficient to maintain a homogenous pluripotent stem cell population. Most likely keeping mESC in suspension in contrast to adherent protocols significantly reduces adhesion mediated integrin signaling, which prevents mESC to differentiate (Hayashi et al., 2007). Our suspension expanded mESC showed a high efficiency of 60% to produce chimeric mice with a high germline transmission rate after more than 3 months in culture (14 passages). This is indeed the gold standard for pluripotency as it shows that the cells contribute to all tissues of the developing animal including the germline.

To quantify the *in vitro* differentiation potential of suspension expanded mESC, we used an established neural tube organoids formation protocol (Meinhardt et al., 2014) and quantified the morphology and transcription factor expression profiles of the cells using imaging and FACS. 1i/LIF suspension cells showed robust lumen formation of organoids after 3 days and neuronal axon formation after 7 days, which is similar to previous characterization of neural tube organoids differentiated from serum/LIF expanded cells (Meinhardt et al., 2014; Ranga et al., 2016).

Finally, while we demonstrated that 1i/LIF suspension culture robustly maintains the pluripotent state of mESC cells over many months, this method also significantly reduced hands-on time and costs compared to standard adherent culture protocols. An earlier comparative study reported 40% less costs of suspension cultures over adherent cultures and less than half the time is spent on maintenance when cells grow in suspension (Tamm et al 2013). However, contrary to this study, using our media composition, cell proliferation remains high in suspension and allows to do more experiments with cells from the running culture as colonies can be collected already after 2 days. Taken together, our results show that 1i/LIF provides an effective and minimal system which maintains pluripotency over the long term, saves time and costs, and enables frequent experimental set-ups. We see this study as a guideline for simplifying mESC culture and hope to encourage more cell biology labs to use stem cell-based organoids as model systems.

## Experimental Procedures

### Maintenance of mouse ES cells

We used mouse R1/E ES cells, a sub-clone from the original R1 ES-cell line, which were derived from mouse crossing 129×1 x 129S1. For suspension culture cells were grown in T25 and T75 flasks with the following media supplements. We used DMEM/F-12+GlutaMAX (Life technology GmbH, cat. no. 31331028) with 1:1 Neurobasal^®^ Medium (Life technology GmbH, cat.no.21103049) supplemented with N2 supplement (Life technology GmbH, cat.no. 17502048), B-27^®^ minus vitamin A (Life technologies, cat. no. 12587010), NEAA (Life technology GmbH, cat.no.11140035), 0.1 mM β-mercaptoethanol (Life technology GmbH, cat.no.21985023). This basal medium was supplemented with <u>2i-LIF</u> (3 µM CHIR99021, Sigma-Aldrich, cat. no. SML1046 and 1 µM PD0325901 Sigma-Aldrich, cat. no. PZ0162 and 1000 U/mL of LIF produced at MPI-CBG, Dresden), <u>1i-LIF</u> (3 µM CHIR99021, Sigma-Aldrich, cat.no. SML1046, cat. no. PZ0162 and 1000 U/mL of LIF) and <u>BMP4-Lif</u> (10 ng/mL of mouse BMP-4 R&D Systems,Inc., cat. no. 5020-BP-010 and 1000 U/mL of LIF).

Suspension cultures were set up in T25 flasks for 3 days and then transferred in T75 with 50% fresh medium. Cells were passaged once a week by filtering spheroids through a 40 µm cell strainer and subsequent washing with PBS. Cells were dislodged from the strainer with 2 ml of Accutase (Life technology GmbH, cat.no. A1110501) into a 10 cm dish. Cell spheres were incubated 2 min at 37 °C and then slowly pipetted (2ml serological pipettes) up and down to make a single cell suspension. After another 2min incubation 6 ml of growth medium were added and cells were again slowly pipetted (10 ml serological pipettes) up and down.

R1 ESC cells in adherent culture were grown in Falcon^®^ 100 mm TC-treated Cell Culture Dish cat.no.353003 in: DMEM (Life technology GmbH, cat. no. 10569010), NEAA (Life technology GmbH, cat. no. 11140035), 0.1mM β-mercaptoethanol (Life Technologies, cat. no. 21985023), 15% FBS (ThermoFisher Scientific cat. no. 10270106), 1000 U/mL of LIF (MPI-CBG, Dresden). Adherent culture was split every 2 or 3 days with medium exchange in between.

### Neuroepithelial cyst formation in Matrigel

Single cell suspensions of ES cells were prepared using Accutase as described above. A suspension of 0.3*10^6^ cells per ml was made in a 70% final dilution of GFR Matrigel (Corning, cat. no. 35623) on ice. Drops of 30 µl Matrigel/Cell solution were seeded into plastic dishes and incubated on ice for 10 minutes. Dishes were then incubated at 37°C for 15 minutes. When the gel was solidified, 2 ml of neuronal differentiation N2B27 medium was added: DMEM/F-12 + GlutaMAX (Life technology GmbH, cat. no. 31331028) with 1:1 Neurobasal^®^ Medium (Life technology GmbH, cat. no. 21103049), supplemented with N2 supplement (Life technology GmbH, cat.no. 17502048), B-27^®^+vitamin A (Life technology GmbH, cat.no.17504044), 500 uM β-mercaptoethanol (Life Technologies, cat.no.21985023), NaPyruvate (Life technology GmbH, cat. no. 11360039) + Glutamate (Life technology GmbH, cat. no. 25030081): Mix 5,5 ml of L-Glutamine and 8,25 ml of sodium pyruvate, store in aliquots at −20°C.

### Proliferation analysis

In suspension culture cells were passaged every 7 days as described above. The total cell number for each flask was evaluated after each passage by counting cells in a Neubauer chamber, excluding dead cells with a Trypan blue staining. For adherent culture the total cell number from each dish was evaluated after passaging every 2 - 3 days by counting cells in a Neubauer chamber, excluding dead cells with a Trypan blue staining. To determine the growth curves the cumulative sum of cell numbers for all passages was calculated. The growth rate P [Cells/Day] was determined from the slope of a linear fit of the growth curves.

### Generation of chimeric mice and determination/quantification of germ line transmission

8 collected ES-Cells were injected into 8-cell C57Bl6/NCrl wildtype embryos. The donor embryos were generated by natural mating. For an efficient ES-cell injection we use the laser assistant injection technology (Poueymirou et al., 2007). About three hours after the ES-Cell injection we transferred 10 - 12 embryos into the oviduct of a pseudogravide Crl:CD1(ICR) female.

To quantify germ line transmission, we analysed the sperm of chimeric male animals. To quantify the amount of donor (R1) and acceptor (C57Bl6/NCrl) derived sperm cells we used short tandem repeats (STR) analysis (GVG Diagnostics, Leipzig, Germany). The STR analysis allows to determine the percentages of ES-cells derived sperms without germline breedings.

### Immunocytochemistry

mESCs cysts were recovered from Matrigel using a cell recovery solution (Corning, cat. no. 354253) by gently pipetting up and down the pre-cooled solution until the Matrigel was dissolved. The suspension was washed in PBS, centrifuged at 1100g for 3 minutes. The pellet was resuspended in Accutase at room temperature, every 5 minutes the solution was gently pipetted up and down, checked on the microscope until most cysts were dissolved to single cells. The suspension was washed in PBS and centrifuged at 1100g for 3 minutes. Cells were fixed in 4% PFA and then permeabilized in cold methanol for 10 minutes at −20 °C. Blocking was done in 5% serum + 0.3% Triton X-100 in PBS. Cells were immuno-stained with (1) rabbit anti-Nanog D2A3 (Cell Signalling, cat. no. 8822), (2) rabbit anti-Sox1 (Cell Signalling, cat. no.4194), (3) rabbit anti-Sox2 D9B8N (Cell Signalling, cat. no. 23064), (4) rabbit anti-Oct4 D6C8T (Cell Signalling, cat. no. 83932) and Dapi in 1% BSA + 0.3% Triton x-100 in PBS.

### Flow cytometry analysis

Flow cytometry was performed on FACSAria system (Becton–Dickinson, Franklin Lakes, NJ, USA). More than 10^4^ cells were analyzed for each sample. Data analysis was performed using FlowJo and MATLAB. The threshold values for gating between ‘ON’ and ‘OFF’ cells were determined by measuring intensity histograms of cells stained with secondary antibody only, for each sample separately. The intensity threshold was set to where the log(intensity) histogram of the secondary only sample dropped to 5% of the maximum peak.

